# Inferring B cell phylogenies from paired heavy and light chain BCR sequences with Dowser

**DOI:** 10.1101/2023.09.29.560187

**Authors:** Cole G. Jensen, Jacob A. Sumner, Steven H. Kleinstein, Kenneth B. Hoehn

## Abstract

Antibodies are vital to human immune responses and are composed of genetically variable heavy and light chains. These structures are initially expressed as B cell receptors (BCRs). BCR diversity is shaped through somatic hypermutation and selection during immune responses. This evolutionary process produces B cell clones, cells that descend from a common ancestor but differ by mutations. Phylogenetic trees inferred from BCR sequences can reconstruct the history of mutations within a clone. Until recently, BCR sequencing technologies separated heavy and light chains, but advancements in single cell sequencing now pair heavy and light chains from individual cells. However, it is unclear how these separate genes should be combined to infer B cell phylogenies. In this study, we investigated strategies for using paired heavy and light chain sequences to build phylogenetic trees. We found incorporating light chains significantly improved tree accuracy and reproducibility across all methods tested. This improvement was greater than the difference between tree building methods and persisted even when mixing bulk and single cell sequencing data. However, we also found that many phylogenetic methods estimated significantly biased branch lengths when some light chains were missing, such as when mixing single cell and bulk BCR data. This bias was eliminated using maximum likelihood methods with separate branch lengths for heavy and light chain gene partitions. Thus, we recommend using maximum likelihood methods with separate heavy and light chain partitions, especially when mixing data types. We implemented these methods in the R package Dowser: https://dowser.readthedocs.io.

## Introduction

The human immune system depends on B cells to produce antibodies that successfully neutralize pathogens. Antibody structures are first expressed as B cell receptors (BCRs), membrane bound immunoglobulins (Ig) on the surface of B cells^1^. A BCR is made up of a heavy chain and a light chain. Both chains are first produced through stochastic V(D)J recombination (only V and J on the light chain) and modified through a second process called affinity maturation^1^. During affinity maturation, B cells undergo rounds of somatic hypermutation (SHM), proliferation, and selection for binding affinity^2^. SHM introduces mutations into the BCR heavy and light chain loci at a rate orders of magnitude higher than that of the background rate of somatic mutations. This evolutionary process produces B cell clones, which are cells that relate by point mutations and indels from a common V(D)J arrangement. Because affinity maturation is a form of evolution by natural selection, techniques from evolutionary biology, such as phylogenetics, can be a powerful means of studying B cells in the context of infection, autoimmunity, and other conditions^3,4^.

While both the BCR heavy and light chains are important for antigen binding, B cell phylogenetic trees have largely been built using only heavy chain information^5^. This was primarily because most bulk BCR repertoire sequencing protocols separated heavy and light chain sequences, so most analyses focus solely on heavy chains due to their greater role in antigen binding and ease of identifying clonally-related sequences due to their higher CDR3 diversity^5–7^. However, recent advances in single cell sequencing now provide paired heavy and light chain sequences for individual B cells^8^. This raises the possibility of building phylogenetic trees using combined heavy and light chain sequences. However, it is unclear how heavy and light chain sequences should be combined in phylogenetic models, what tree building methods should be used to incorporate paired data, and whether this significantly improves B cell phylogenetic trees. Including light chains has shown mixed improvements in B cell clonal identification so it is plausible that heavy chains alone will sufficiently resolve most B cell trees^7,9^.

Here, we investigate which methods estimate the most accurate phylogenetic trees from paired heavy and light chain BCR sequences, and test whether they improve tree accuracy over using heavy chain sequences alone. We implement these methods in Dowser v2.0.0, an R package that is part of the Immcantation framework, which provides tools for B cell phylogenetic analysis and visualization^10^. Using simulated and empirical datasets, we show that paired heavy and light chain sequences significantly improve accuracy and reproducibility of B cell phylogenetic trees. This improvement was consistent across all tree building methods. Crucially, the worst-performing method using paired heavy and light chain sequences had a higher accuracy than the best-performing method using only heavy chain sequences. Results were mixed, however, when some cells were missing light chains. Light chains can be missing due to several reasons, such as drop-out during single cell sequencing, or from mixing single cell data with bulk BCR heavy chain sequencing. We show that when only some BCRs have paired heavy and light chains, most commonly used phylogenetic tree building methods produce significantly biased branch length estimates. We also show that maximum likelihood methods with multiple partitions, in which branch lengths are estimated separately for heavy and light chain sequences, have superior performance when only some cells have paired heavy and light chains, such as when using mixed single cell and bulk BCR data.

## Methods

### Building B cell phylogenetic trees with paired heavy and light chain sequences with Dowser

Phylogenetic tree building methods require sequences to be orthologous and in alignment. Dowser takes multiple steps to ensure both heavy and light chain sequences are properly aligned before building trees. Once separated into clones, this is straightforward with heavy chain sequences. However, depending on how clonal clustering is performed, B cell clones can contain cells with distinct light chain rearrangements that do not descend from the same light chain VJ rearrangement. Further, individual B cells can express multiple distinct light chains^11^. To resolve ambiguities in light chain sequences within a clone, the following steps are taken in Dowser after first grouping B cells into clones:

1. For all cells within a clone, if a heavy chain is paired with multiple or ambiguous light chain rearrangements, enumerate all possible combinations of ambiguous V and J genes.
2. Group cells based on light chain V and J gene assignment and junction lengths, starting with the largest possible grouping. Resolve ties by assigning to the subgroup with the lowest index.
3. If a heavy chain has no paired light chain, assign that cell to the same subgroup as the most similar heavy chain sequence within the same clone, determined by the Hamming distance. Resolve ties by assigning them to the largest subgroup. Resolve remaining ties by assigning them to the subgroup with the lowest index.
4. Remove light chains with no paired heavy chain.
5. Treat separate V and J subgroups as separate clones.
6. Reconstruct clonal V and J germlines for each chain.

Steps 1-4 are implemented in the Dowser function *resolveLightChains*, step 5 is implemented in the function *formatClones*, and step 6 is implemented in the function *createGermlines*. Within the Dowser workflow, each cell must have exactly one heavy chain sequence, and both heavy and light chain sequences must be in IMGT-gapped alignment. Performing steps 1-6 ensures that all cells within each clonal group have exactly one heavy, at most one light chain, and that all sequences within a clone are in alignment with their predicted common ancestor.

To create a combined alignment, each paired heavy and light chain sequence is concatenated together. Inferred heavy and light germline sequences within a clonal subgroup are similarly concatenated and used as the root of the tree. Missing light chains are represented as a string of ambiguous N nucleotides of the same length as the other sequences in the alignment. Concatenated sequences that are identical or differ only by ambiguous characters are collapsed into one sequence by default.

Dowser supports building trees with six external programs whose outputs are analyzed within a common framework (**Table 1**). To test which method performed the best, we used all six programs in testing and included two additional models with multiple gene partitions (hereafter “multi-partition models”) for a total of 8 tree building methods. Two methods were based around maximum parsimony: PHYLIP’s *dnapars* v3.697 and the R package *phangorn*’s v2.10.0 maximum parsimony function ‘pratchet’^12,13^. For *dnapars*, rearrangements were searched on one best tree topology^12^. In *phangorn*, default parameters of the “pratchet” function were used. Four methods were based on maximum likelihood using a single-partition for both heavy and light chains: PHYLIP’s *dnaml* v3.697, *phangorn*’s maximum likelihood function ‘pml,’ IgPhyML v2.0.0, and RAxML v1.2.0^12–15^. Default search parameters were used in *dnaml*, and for maximum likelihood estimation with *phangorn*, tree topology, branch lengths, and model parameters were estimated using the general time reversible (GTR) model. With IgPhyML and RAxML, two different models were tested: one single-partition model and a multi-partition model. The multi-partition models were “scaled” models, which allowed heavy and light chain branch lengths to differ proportionally by a scalar factor estimated by maximum likelihood. In both RAxML options, the GTR model was used, and parameters were estimated for each clone and partition separately. For IgPhyML, the HLP19 model was used, and parameters were shared among all clones and partitions within a repertoire^14^. Within Dowser, all of these tree building methods can be specified using different options of the *getTrees* function.

**Table 1.**
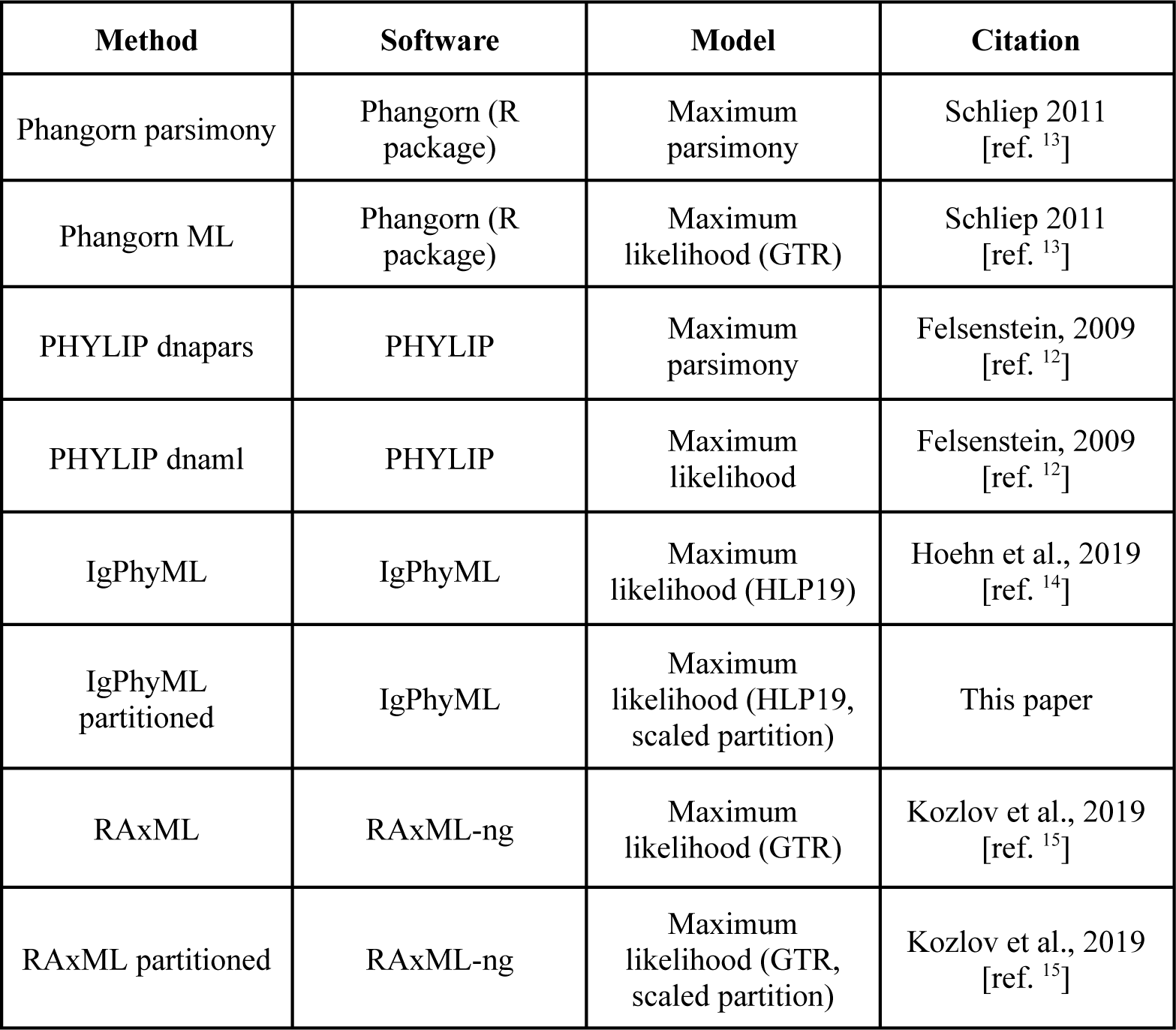
Tree building methods tested. All are supported by Dowser v2.0.0.

### Simulations

We simulated paired affinity maturation of heavy and light chain sequences using a combination of two methods: the *bcr-phylo* simulation program and the R package *SHazaM* v1.1.2^16,17^. *Bcr-phylo* simulates lineages of B cells undergoing affinity maturation over a specified number of generations. To obtain biologically realistic paired heavy and light chains as starting sequences, we used previously published empirical single cell dataset of naive B cells (see **Empirical datasets**)^18^. Simulations begin with a randomly chosen heavy chain from this dataset. For each of the 20 repetitions, we simulated 50 clones with 50 sequences each. Initial analyses showed that simulations over 125 generations produced sequences with similar heavy chain SHM as in two empirical dataset (**Fig. 1**). Thus, all simulations were performed for 125 generations. Lineages were simulated with selection based on amino acid sequence similarity to a pre-specified target sequence. Other parameters were kept as default. The resulting trees were used as the true trees for benchmarking.

**Figure 1.**
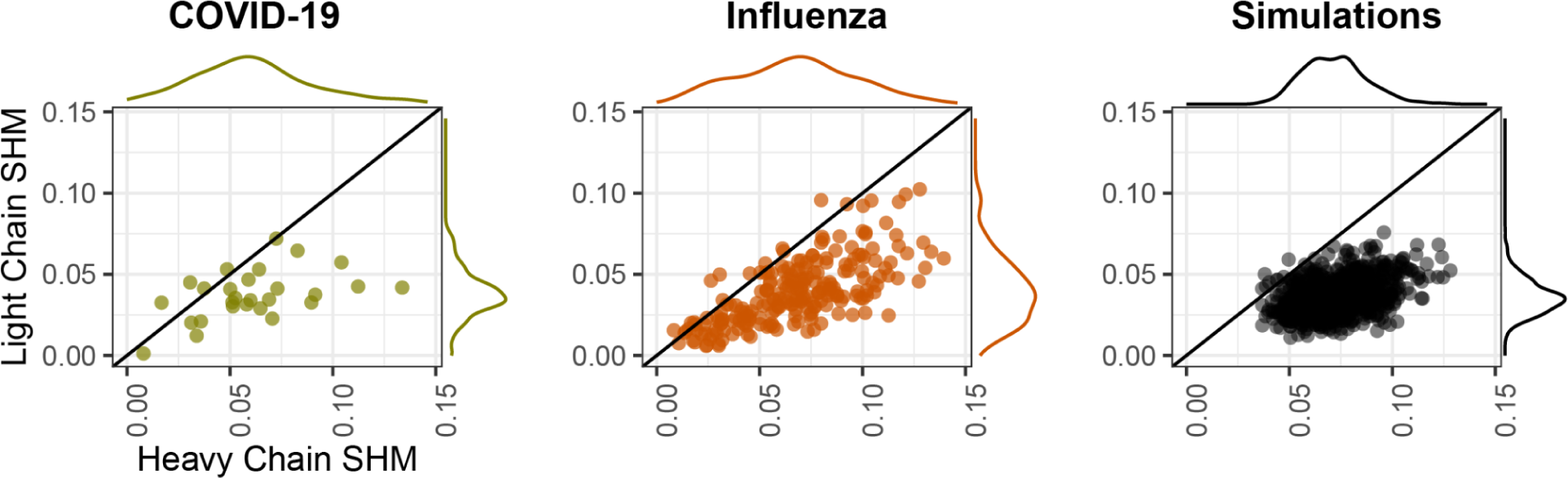
Comparison of heavy and light chain somatic hypermutation (SHM) among datasets. Each point shows the mean SHM frequency along the heavy and light chain V-gene for each clone. This is shown for a dataset of subjects with COVID-19 (left), a subject who recently received an influenza vaccination (middle), and 20 simulated datasets (right). For empirical datasets, only clones with at least three or five (COVID-19 and influenza datasets, respectively) distinct sequences were included. The line partitioning each plot indicates equally mutated heavy and light chains. Marginal distributions for heavy or light chain SHM are shown outside of each plot.

Initial mutational frequency analysis of the two empirical datasets showed light chains have approximately half the SHM frequency of heavy chains (**Fig. 1**), and previous work has shown light chains have different SHM hot- and cold-spots from heavy chains^19^. However, *bcr-phylo* does not currently allow paired light chains to evolve at a different rate from paired heavy chains. To simulate paired chains with different mutation rates, we used the lineage trees generated by *bcr-phylo* but simulated the mutations in heavy and light chains on each branch separately using *SHazaM*. For heavy chains, we iteratively mutated the starting sequence using the S5F model of heavy chain SHM hot- and cold-spots^20^. Starting at the germline node, the number of mutations was drawn from a Poisson distribution with a mean set as the branch length between each node and its descendants. This process repeated until all tip sequences were simulated. Light chains were simulated using a similar process, but the HKL_S5F model of light SHM chain hot- and cold-spots was used, and branch lengths were halved to account for observed differences in light chain SHM (**Fig. 1**)^19^. For light chain starting sequences, we used the light chain paired with the starting heavy chain sequence in the empirical naive B cell dataset. In summary, heavy and light chains within each clone were simulated using the same tree topology, but differing mutation processes.

To compare branch length estimation among different tree building methods, we also performed simulations using simple triplet trees. These trees consisted of a germline, two tips, and three branches. For these simulations, heavy and light chains were randomly chosen from the same naive B cell dataset as other simulations (**Empirical datasets**). Along each branch, 25 mutations were added to the heavy chain according to the S5F model, and 12 mutations were added to the light chain according to the HKL_S5F model. This process was repeated for 1000 repetitions. We made two versions of the simulated datasets. In the single cell only (SC) data, both heavy and light chains of all tips were retained. In the mixed single cell and heavy chain BCR bulk (SC+bulk) data, we removed the light chain of one tip. We then estimated branch lengths from all simulated data and compared them to the true tree lengths. These simulations and analyses used the same packages as others, except R 4.2.2, *SHazaM* v1.1.2, RAxML-ng v 1.1.0, and *alakazam* v1.2.1.999^21,17,15^.

### Empirical BCR datasets and processing

Single cell datasets of B cells were obtained from three published sources. The first was a study of hospitalized and non-hospitalized COVID-19 patients with healthy controls, and was used to generate a dataset of naive B cells for simulations^22^. Naive B cells were identified based on 1) prior assignment as naive B cells based on scRNAseq data in the source publication, 2) IGHM or IGHD constant region, and 3) unmutated heavy chain BCR. To determine the level of heavy chain SHM in this dataset, we first identified clones by grouping cells within a subject based on common IGHV genes, IGHJ genes, and junction lengths. We then used single-linkage hierarchical clustering to group cells with at least 80% junction similarity using *SCOPer*^23^. Within these clones, we reconstructed consensus germline sequences using the *createGermlines* function in Dowser v2.0.0^10^. SHM was quantified as the Hamming distance from these germline sequences along the V gene (IMGT positions 1-312) for each heavy and light chain sequence in the dataset^17^.

The second empirical dataset was a study of 10 hospitalized COVID-19 patients sampled at two timepoints^18,22^. Clonal clustering was performed in the original study. Only clones with at least three unique paired heavy and light chain sequences were included. This resulted in 27 clones ranging in size from 3 to 61 unique paired heavy and light chain sequences (mean 6.89).

The third dataset was a study on influenza vaccination that investigated a single subject using peripheral blood and lymph node samples obtained at multiple timepoints after vaccination^24^. This dataset included both single cell BCR and bulk heavy chain BCR sequences. More specifically, there were 101,048 paired single cell BCR sequences and 43,751 bulk heavy chain BCR sequences. Lymph node sequences were derived only from single cell sequencing data, while blood sequences were from a mixture of bulk (48.40%) and single cell (51.60%) data. For some analyses, only single cell data was included (e.g. **Fig 1**), and in others both bulk and single cell data were used. Clonal clustering was performed as part of the original study. When using only single cell BCR data, only clones with at least 5 unique paired heavy and light chain sequences were included. This resulted in 227 clones varying in size from 5 to 41 paired sequences (mean 8.01). The full dataset of mixed single cell and bulk BCR data also included sequences of monoclonal antibodies that were experimentally verified to bind to influenza. When analyzing this full dataset, only clones containing a monoclonal antibody were included. Further, only clones with at least three unique heavy and light chain sequences, as well as at least one day 5 blood sequence and at least one germinal center B cell were included. This resulted in 38 clones ranging in size from 3 to 309 unique sequences (mean 36.24).

Due to their high frequencies of expanded clones, the second and third datasets were used as comparisons to simulated data (**Figs. 1-3**). B cells with multiple heavy chains were removed. B cell clones with multiple light chain rearrangements (V and J genes) were split using Dowser v2.0.0^10^. For each locus in each clone, unmutated germlines sequences were constructed based on alignments to germline V and J reference sequences in the IMGT database^25^. Within heavy chain germlines, D segments and N/P regions were masked using ‘N’ nucleotides. As in the first data set, somatic hypermutation was quantified as the length-normalized Hamming distance from its inferred germline along the V gene region (IMGT positions 1-312).

### Evaluation metrics

To quantify the accuracy of tree topology, we calculated the Robinson-Foulds (RF) cluster distance between an estimated tree and its true topology^26,27^. Because ground truth trees are not available in empirical BCR data, this was only possible in simulations. The RF distance is widely used to compare trees and is equal to the number of unique bipartitions between two trees. The RF cluster distance is a modification, in which branches with very small lengths are collapsed into polytomies.^27^ We collapsed branches that were shorter than 0.001 mutations per nucleotide or 0.003 mutations per codon. The RF cluster distance was then calculated as the number of subclades unique to each tree. We used Dowser’s calcRF distance function for these calculations, which does not require trees to be strictly binary. This approach is based on the RF cluster distance in Zhang et al.^27^ Because we calculate the RF cluster distance from the true tree, a lower distance indicates a more accurately estimated topology^26^.

In the absence of a ground truth tree, reproducibility is often used as a metric of support for particular clades, and is typically measured using phylogenetic bootstrapping^28,29^. After building a tree (called the “full tree”) using all sequence sites, we randomly sampled codon sites with replacement and then re-built the tree using this resampled data. We repeated this 100 times, creating bootstrap replicate trees. The bootstrap score for each branch in the full tree was then calculated as the number of times that branch, defined by its subtaxa, was found in the 100 bootstrap replicate trees. The higher the score, the more reproducible the branch. We then calculated the mean bootstrap score across all branches within the full tree. Because bootstrapping does not require a ground truth, we calculated bootstrap scores for both simulated and empirical data. This was performed using the Dowser v2.0.0 function *getBootstraps*.

While RF cluster distance and bootstrapping focus purely on tree topology, branch length estimates are another important part of phylogenetic trees, and represent the estimated number of mutations per site (codon or nucleotide). Because we focus on large scale trends rather than particular branches, we report the tree length, which is the sum of all branch lengths within a tree, and represents the total number of mutations that occurred per site within the lineage. When a ground truth is available through simulations, we report the percent error in the estimated tree length, which is the true length minus the estimated length divided by the true length.

In all analyses in which trees built with paired heavy and light chains are compared to trees built with only heavy chains, sequences used for heavy chain-only trees were not collapsed but were instead filtered to include only the cells remaining in the heavy and light chain dataset after collapsing. This ensured that heavy chain-only trees were built using sequences from the same B cells as with paired heavy and light chain trees, and that the resulting trees could be appropriately compared using the above metrics.

## Results

### Inferring and simulating paired heavy and light chain affinity maturation

Inferring B cell phylogenies from paired heavy and light chain sequences is similar to species tree inference, in which a single tree is inferred using multiple genes^30^. Some aspects of B cell biology make phylogenetic inference using paired heavy and light chains less complicated than in other systems. For example, because B cells reproduce by binary cell division, heavy and light chain sequences within a clone have the same underlying evolutionary history, and processes such as incomplete lineage sorting should not occur. Because of this linked evolutionary history, we concatenate heavy and light chain sequences within each clone and treat them as a single alignment. However, because these genes have distinct roles and are located on different chromosomes, it is possible they experience different mutation rates and/or selection pressures^1^. It is important to quantify these differences and to design methods that take them into account.

There are multiple ways heavy and light chains could experience different mutation and selection patterns. Prior work using heavy chain bulk BCR sequences showed that heavy and light chains have distinct somatic hypermutation (SHM) hot- and cold-spots and that, on average, light chain sequences have approximately half the frequency of SHM as heavy chain sequences^19^. To demonstrate this latter result was consistent when using paired heavy and light chains, we compared SHM frequency in two published single cell sequencing datasets: one obtained from 10 hospitalized COVID-19 patients and another from a subject recently vaccinated against influenza^22,24^. In both datasets, light chain sequences had approximately half the number of mutations per site as their paired heavy chain in the same cell (**Fig. 1**). This indicates that light chains evolve at roughly half the rate of the heavy chains.

There are many available tree building methods, which may differ in their appropriateness for paired heavy and light chain trees. Maximum parsimony approaches estimate the tree topology and set of branch lengths that minimize the number of mutations that occurred within a tree. This is an intuitive and popular approach for B cell lineage trees but may be less accurate for trees with longer branches^27^. Maximum likelihood approaches use a Markov model of sequence evolution to estimate the tree topology and set of branch lengths that maximize the likelihood of a set of sequence data. In standard, single-partition maximum likelihood approaches, both heavy and light chain sequences are treated as part of the same gene, and a common tree topology, set of branch lengths, and model parameters are estimated. In maximum likelihood models with multiple gene partitions (multi-partition models), some features can be estimated separately for heavy and light chain sequences. Here, we test “scaled” multi-partition models in which heavy and light chains share a common tree topology, but heavy and light chain branches are proportionally shrunk or expanded by a scaling factor. In one method (RAxML), all model parameters are estimated separately for each partition in each clone, and in another (IgPhyML v2.0.0), a common set of model parameters and scaling factors is shared among all clones within a repertoire. To determine which method performed best for heavy and light chain sequences, we tested eight tree building methods, including two using maximum parsimony, four using single-partition maximum likelihood, and two using multi-partition maximum likelihood (**Table 1**). These are all supported within a common tree building and visualization framework in Dowser v2.0.0.

Testing the accuracy of B cell lineage trees requires a ground truth. Because this is not currently available for empirical B cell lineages, we performed simulations representing affinity maturation (**Methods**). Briefly, we first generated trees by simulating affinity maturation of heavy chain sequences using the simulation package *bcr-phylo*^16^. To simulate paired heavy and light chain evolution, we then 1) selected paired heavy and light chain starting sequences from a dataset of naive B cells, 2) placed them at the germline position of the tree topology generated by *bcr-phylo,* 3) iteratively added mutations according to the branch lengths of the tree until all tip sequences were simulated. Only sequences at the tips were retained. Mutations were added according to published models of heavy and light chain SHM hot- and cold-spots^19,20^. Further, light chain mutations were added at half the rate of heavy chain mutations. We repeated this process for 20 replicates. We then confirmed these simulations produced heavy and light chain SHM frequencies similar to two empirical single cell datasets (**Fig. 1**).

### Incorporating paired light chains improves tree accuracy and reproducibility

A primary goal of phylogenetics is to reconstruct ancestor/descendant relationships within a lineage. This is reflected in a phylogenetic tree’s topology. To quantify how accurately a tree’s topology has been estimated, we calculate the Robinson-Fould’s (RF) cluster distance between a B cell clone’s estimated and true phylogenetic tree topology (**Methods**)^27^. Lower RF cluster distances, in this case, indicate a more accurate topology^26^. We find that in all eight tree reconstruction methods, building trees using paired heavy and light chain sequences had significantly lower average RF cluster distances from the true tree compared to only using the heavy chain (**Fig. 2**, **Supplementary Fig. 1A**). This indicates that in these simulations, trees constructed from paired heavy and light chain sequences are more accurate than those constructed from heavy chains alone.

**Figure 2.**
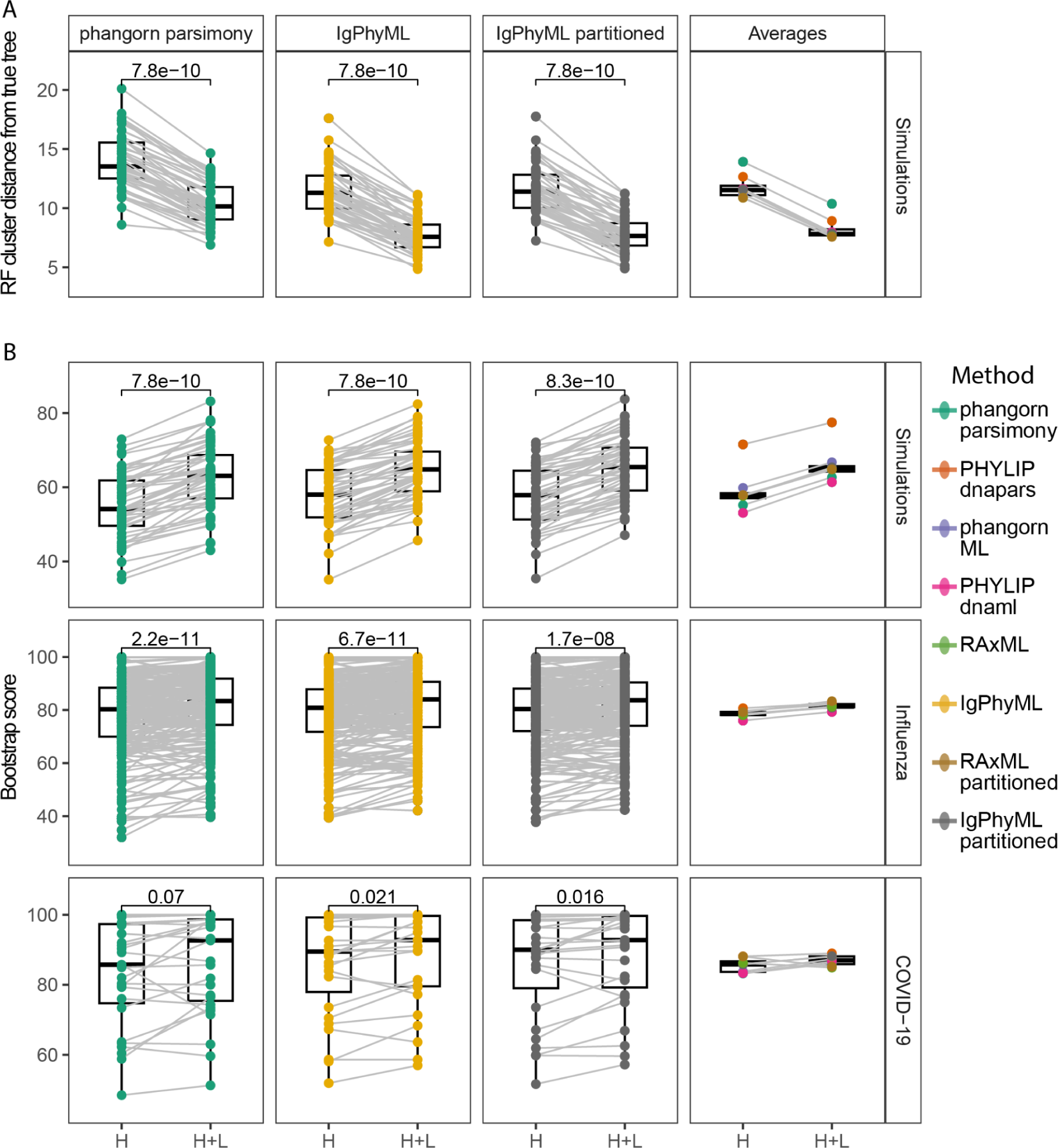
The accuracy and reproducibility of tree topology estimates are improved using paired heavy and light chains. A) Robinson-Foulds (RF) cluster distance between estimated and true tree topologies for trees built using only the heavy chain (H) and paired heavy and light chain sequences (H+L) using a representative maximum parsimony, single-partition maximum likelihood, and multi-partition models. Additionally, the right column shows the average RF distance for all methods. B) Mean bootstrap values for trees built using each dataset. P values were calculated using a Wilcoxon test. For comparisons with all methods, see **Supplementary Fig. 1**.

Improvement in accuracy from including light chains was greater than that of different tree building methods using only the heavy chain. More specifically, among all tree building methods, using paired heavy and light chains improved mean RF cluster distance by an average of 3.55, while the range between methods using only heavy chains was only 3.04 (**Fig. 2A**). On average, the worst performing method using paired heavy and light chains (mean RF = 10.37) performed better than the best performing method using only heavy chains (mean RF = 10.88, **Fig. 2A**). This indicates that, among the simulations and methods tested, the decision to include the light chain is likely more important than the choice in tree building method.

Another important aspect of phylogenetic trees is reproducibility, which is used as a measure of support for particular clades. We measure this using phylogenetic bootstrapping, which is performed by randomly sampling sequence alignment sites with replacement, re-building the tree, and quantifying whether the same clades in the tree are reproduced^28,31^. This is repeated for 100 bootstrap replicates, resulting in a bootstrap score for each clade that indicates the number of replicates containing the same clade. A higher bootstrap score indicates a more reproducible branch, with a score between 70-90 generally considered reliable^29^. Using the same simulation data as above, we found that incorporating light chain sequences produced trees with significantly higher average bootstrap scores compared with using only heavy chain sequences, irrespective of the tree building method (**Fig. 2B**). Because bootstrapping does not require a ground truth, we tested whether incorporating light chain sequences increased the reproducibility of trees in our empirical influenza and COVID-19 datasets. In all methods tested, incorporating light chain sequences significantly improved the reproducibility of trees built in the empirical influenza dataset (**Fig. 2B**, **Supplementary Fig. 1**). By contrast, only four of the eight methods gave significantly higher bootstrap scores in the COVID-19 dataset, perhaps due to its fewer large clones compared to the influenza dataset (**Supplementary Fig. 1**).

### Partitioned maximum likelihood methods have superior performance in mixed bulk and single cell BCR data

Single cell sequencing of BCRs currently has significantly lower throughput than bulk BCR sequencing methods. Because of this, single cell and bulk heavy chain BCR sequencing from the same sample are often combined for higher resolution of B cell clones^32,33^. While this can be advantageous, it also creates a non-random missing data problem in which some heavy chains have paired light chains while others do not. Light chain data can also be missing when there is drop out in single cell sequencing. Tree building methods differ in how they deal with missing data and, thus, likely differ in their ability to handle missing light chains.

To determine how trees should be built when only some heavy chains have paired light chains, we re-analyzed the simulations in **Fig. 2** after randomly discarding light chains from different proportions of B cells. Sequences were discarded such that all clones had at least one cell with a paired light chain. We found that across all methods, the accuracy of estimated tree topologies generally decreased as more light chains were removed, reaching approximately the same accuracy and as trees built with only heavy chains if ≥30% of cells had missing light chains (**Fig. 3A**). Interestingly, some maximum likelihood methods performed slightly better when nearly all 80-95% of light chains were removed than when there were an even number of paired and missing light chains (**Fig. 3A**). In general, maximum likelihood methods, especially RAxML, performed better than maximum parsimony methods in these benchmarks. However, most methods performed competitively, within a range of 2-3 more edits from the true tree on average (**Fig 3A**). In terms of the estimated tree topology, multi-partition models did not perform significantly better than their single-partition counterparts (**Fig. 3A**). Overall, these results indicate that tree topology can still be improved in most methods even if only a portion of cells have associated light chains.

**Figure 3.**
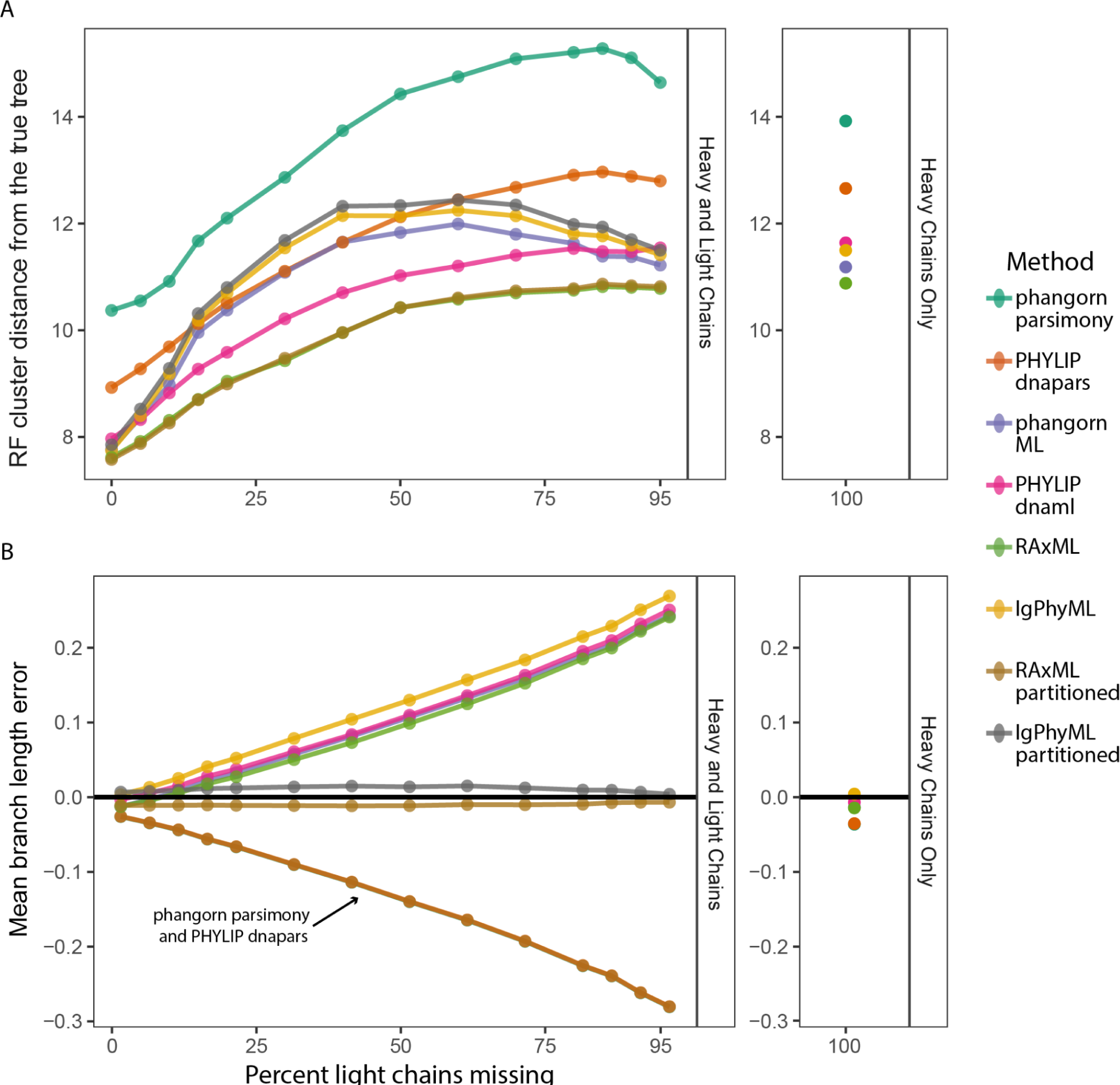
Effect of missing light chains on tree topology and branch length estimates. A) Average RF cluster distance for trees built using paired heavy and light chain sequences (H+L, left) with an incrementally greater proportion of light chains missing. The top right panel shows the mean RF distance for trees built using only the heavy chain (H). B) Mean branch length error of trees built using H+L sequences (left) and H sequences (right). Points show the means of all 20 simulation replicates.

Branch lengths are another important aspect of phylogenetic trees and typically represent the estimated number of mutations per site. Using the same simulated data as above, we compared the estimated tree length, which is the sum of all branch lengths within the estimated tree, to the true tree length for each clone. Methods differed strikingly in their performance and separated into groups based on fundamental model frameworks (**Fig. 3B**). When some light chains were missing, both maximum parsimony methods substantially underestimated tree length. This effect worsened as more light chains were removed, reaching nearly 30% error when 95% of cells were missing light chains. Maximum parsimony methods also tended to underestimate tree length when using only heavy chains, though to a much lesser extent (**Fig. 3B**). Single-partition maximum likelihood methods followed the opposite trend by overestimating tree length as more light chains were missing, reaching >20% error when 95% of light chains were missing. By contrast, multi-partition models were not significantly affected by missing light chains and gave accurate branch lengths (between −1.2 and 1.5% error) in all cases tested. These large differences in performance are explainable by how each method deals with missing data. Maximum parsimony methods typically assume no mutations occurred in missing nucleotides, which would cause them to underestimate mutations when some light chains are missing. Maximum likelihood methods assume data are missing randomly within a partition. Single-partition models likely overestimate branch lengths because they effectively assume mutations in missing light chain sequences occurred at the same rate as their paired heavy chains. In reality, light chains have approximately half the mutations as their paired heavy chains. Multi-partition models, by contrast, allow light chain partitions to have shorter branch lengths than heavy chain partitions and more accurately estimate the number of mutations for missing light chains.

To investigate this effect further, we performed simulations using simple 3-taxa trees. Sequences were simulated down three branches, starting from the germline, with each branch having 25 mutations on the heavy chain and 12 mutations on the light chain. For “single cell” (SC) simulations, paired heavy and light chain sequences were retained for all three tips. To recreate a mixture of single-cell and bulk data, the light chain of one tip was removed. We simulated 1000 datasets and then benchmarked each phylogenetic method’s ability to estimate the length of each branch. All methods accurately estimated branch lengths when heavy and light chains were available for all tips (**Fig. 4**). However, there were significant differences among methods when some light chains were missing, as occurs when using mixed single cell and bulk BCR data. Maximum parsimony methods significantly underestimated the length of ancestral branches to cells with a missing light chain. Interestingly, the ACCTRAN branch length estimation method in *phangorn* also produced multi-modal estimates of the other two branch lengths^13^. All four standard single-partition maximum likelihood methods significantly overestimated the branch leading to the cell with the missing light chain. By contrast, multi-partition models accurately estimated all three branch lengths.

**Figure 4.**
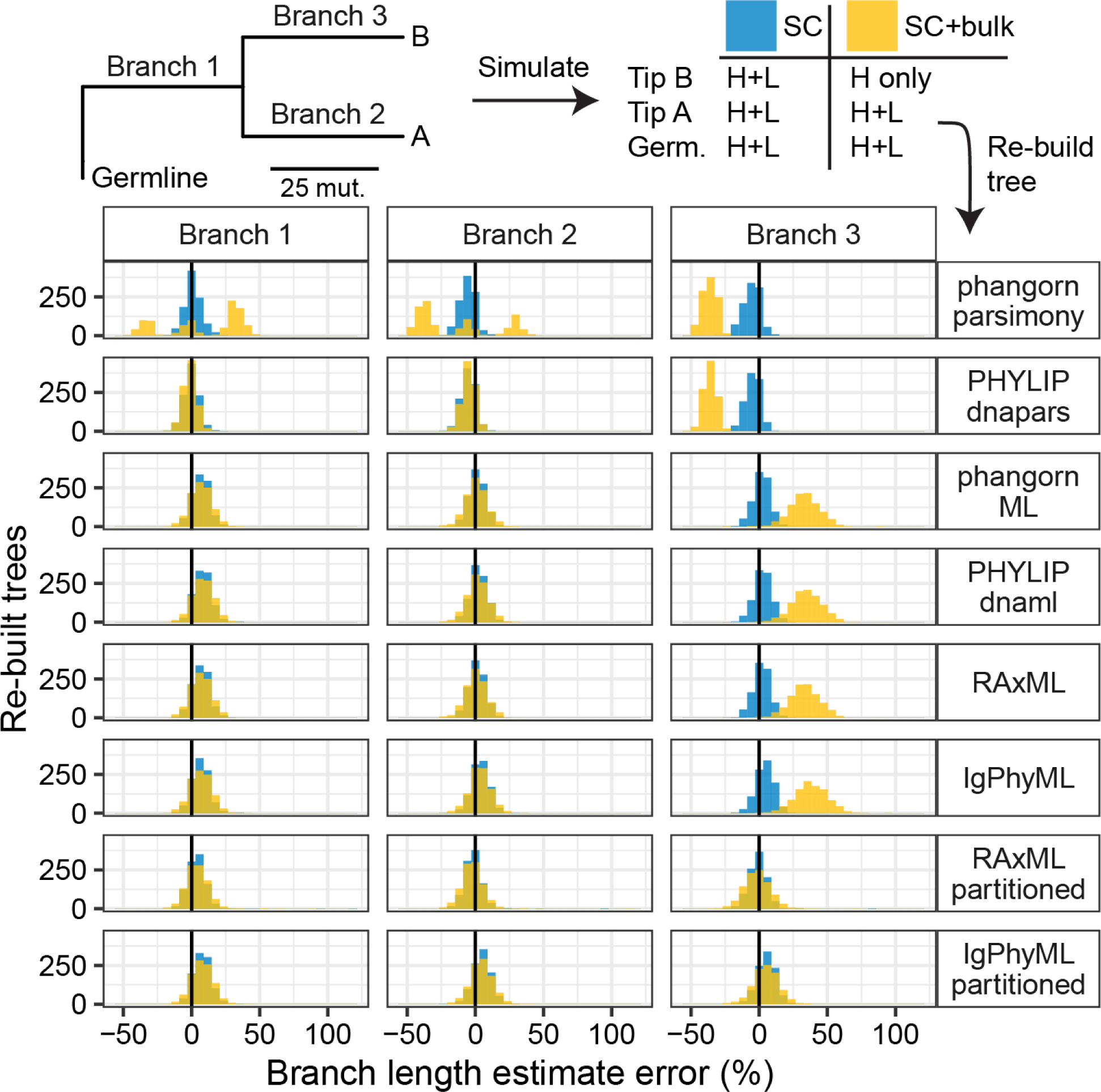
Branch length estimates from many methods are affected by missing light chains. A) Schematic of simulation strategy. Heavy chains were simulated with approximately twice as many mutations as light chains. For single cell (SC) data, heavy and light (H+L) sequences were retained for all three tips. For mixed single cell and bulk (SC+bulk) data, the light chain of tip B was removed. B) For each replicate, branch lengths were estimated using different methods and compared to the true branch length. In SC+bulk data, most methods incorrectly estimated the length of branch 3, which immediately precedes tip B.

Within both simulation experiments performed, multi-partition maximum likelihood methods were the only approach tested that accurately estimated branch lengths when some light chains were missing (**Figs. 3B and 4**). By separating heavy and light chain sequences into different partitions, these methods correctly estimate phylogenetic branch lengths even if light chains were missing for some sequences. These models are available in RAxML, and we implemented a repertoire-wide partitioned model in IgPhyML v2.0.0^14,15^. We note, however, that the benefit of including light chain information diminishes once >30-40% of B cells come from bulk data (**Fig. 3A**). In these cases, using single-partition maximum likelihood methods on heavy chain sequences may give a similar performance, but will not include mutational information from paired light chains.

### Tree-building methods lead to different conclusions about influenza vaccine response

We next tested whether tree building methods could lead to different conclusions in practice by investigating B cell responses to influenza vaccination in humans. Influenza vaccination in humans classically stimulates the expansion of highly mutated plasmablasts in the blood around one week post-vaccination^34^. Recent work has shown that members of these B cell lineages also re-enter germinal centers (GCs) and accumulate new mutations^24,35^. However, it is unclear how GC sequences relate to circulating B cells in the blood. One simple hypothesis is that early GC B cells are as mutated as their clonally related B cells in the blood, but it is possible they are seeded by clonal members with fewer or more mutations.

To test this hypothesis, we used single cell and bulk BCR data from a previously published study using human blood and lymph node samples following influenza vaccination^24^. This dataset included blood samples taken 5 days after influenza vaccination and lymph node samples taken at 5, 12, 28, and 60 days after vaccination. Importantly, all lymph node BCR sequences were obtained from single cell sequencing, while blood BCR sequences were 51.60% single cell data, and 48.40% bulk data. Within lymph node samples, GC B cells were identified in the original study based on paired RNAseq data. This dataset also included sequences of experimentally-verified influenza binding monoclonal antibodies (mAbs), which allow for identification of influenza binding clones. Within this dataset, we identified 38 clones containing influenza-binding mAb sequences, blood BCR sequences, and GC B cells. To simplify interpretation, mAb sequences were removed from the clones before analysis. We then built trees for these clones using *phangorn’s* maximum parsimony, IgPhyML, and multi-partition IgPhyML as representative programs for maximum parsimony, single-partition maximum likelihood, and multi-partition maximum likelihood.

To test if tree building methods could lead to different conclusions, we estimated the divergence – defined as the sum of branch lengths from each tip to the germline – for all sequences in each clone. This is essentially a tree-based estimate of SHM and is an intuitive way of quantifying the relationship between these two compartments within trees. To test if early GC B cells were more or less mutated as their clonal relatives in the blood, we compared the mean divergence of blood sequences at day 5 to their clonally-related GC sequences at day 12. Then, to understand how mutations accumulated over time in the GC, we investigated GC B cells at day 60, and across all timepoints pooled together (**Fig. 5**). Using maximum parsimony, one would conclude that GC B cells at day 12 were equally as diverged as their clonal relatives in the blood at day 5, and then accumulated significantly more mutations by day 60 (**Fig. 5**). By contrast, using either single or multi-partition maximum likelihood models, one would conclude that GC B cells at day 12 were less diverged than their clonally-related day 5 cells in the blood, but then reached similar levels of divergence by day 60. Single and multi-partition models gave qualitatively similar conclusions, but differences between blood and GC B cells were slightly more pronounced with single-partition models. These conflicting results are explainable by the simulation results in **Figs. 3** and **4**. Because many blood sequences were missing light chains, maximum parsimony likely underestimated the number of mutations in blood B cells. This made them appear to be as diverged as day 12 GC B cells and significantly less diverged than day 60 GC B cells. Because the number of GC B cells were limited at individual timepoints, we next tested whether different tree building methods produced different patterns when all GC timepoints were included. When all GC timepoints were included, maximum parsimony predicted GC B cells were significantly more diverged than day 5 clonal relatives in the blood. Single-partition models predicted GC B cells were significantly less diverged than day 5 clonal relatives in the blood and multi-partition models predicted similar divergences between these cell types. Thus, the choice in tree building methodology significantly impacts the biological conclusion from this analysis.

**Figure 5.**
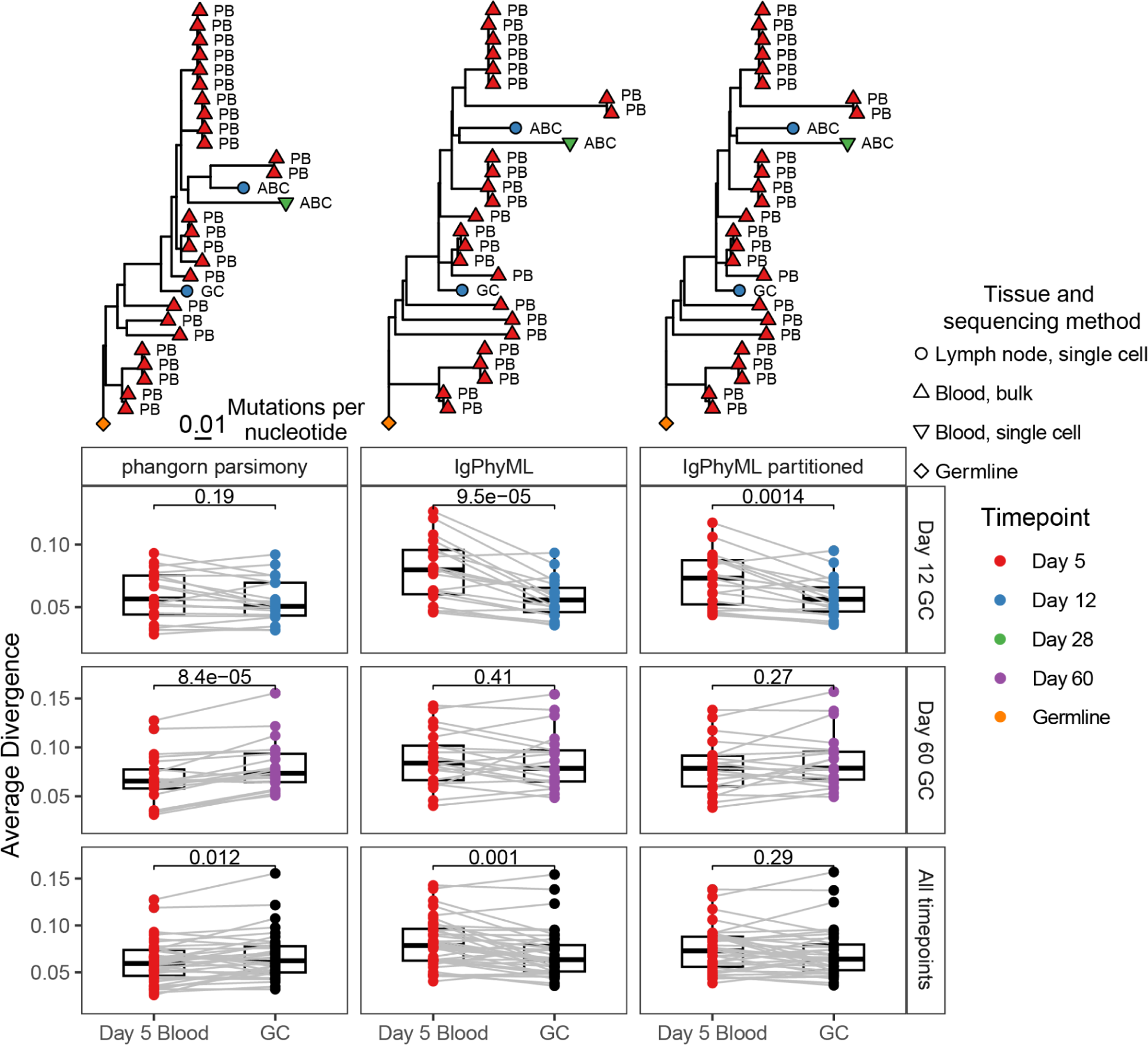
Choice in tree building method significantly impacts conclusions with mixed single cell and bulk data. Data were obtained from blood (48.40% bulk BCR, 51.60% single cell) and germinal center (GC, 100% single cell). Mean divergence (sum of branch lengths from germline to each tip) for blood and GC sequences in shared clones estimated using different methods are shown. Only data from clones sampled in both the GC at the specified time point as well as blood at day 5, are shown. Above each plot is a representative tree. For heavy chain only comparisons, see **Supplementary Fig. 2**.

While a ground truth was not available in this case, we repeated our analyses using only heavy chain sequences and re-built trees with maximum parsimony and single-partition maximum likelihood (**Supplementary Fig. 2**). Using only heavy chains is likely underpowered relative to using paired data (**Fig. 3A**), but is likely less biased due to the lack of missing light chain data (**Fig. 3B**). Both methods predicted significantly lower divergence in day 12 GC B cells compared to clonally-related day 5 blood B cells, consistent with single- and multi-partition maximum likelihood results from paired data (**Fig. 5**). At day 60, maximum likelihood predicted GC B cells were non-significantly more diverged than day 5 blood B cells, while maximum parsimony predicted GC B cells were significantly more diverged. These results are consistent with their respective methods when using paired data (**Fig. 5**). Considering all GC timepoints, neither method found significant difference in divergence between GC B cells and clonally-related day 5 blood B cells, which was consistent with only multi-partition models using paired data (**Fig. 5**). Thus, the tree building paradigms tested produce qualitatively different conclusions, and when using paired heavy and light chain data multi-partition models were the most consistent with results from using only the heavy chain. Because of these results coupled with the simulation results, we recommend using the multi-partition maximum likelihood model when using data with missing light chains. Biologically, these results indicate that when clones contain both GC B cells and blood B cells, early GC B cells tend to arise from less-mutated clonal members, while clonal relatives in the blood arise from more-mutated clonal members. Over time, GC B cells then accumulate slightly, but not significantly, more mutations than their related day 5 clonal relatives in the blood.

## Discussion

Recent advances in single cell sequencing technology now provide paired heavy and light chain BCR sequences for individual B cells. While building trees using paired heavy and light chain sequences has been done in practice, it is not known which phylogenetic methods are most appropriate for building trees with paired heavy and light chains, nor has it been shown whether trees built with paired data are more accurate^36^. While single cell sequencing can provide paired heavy and light chain data, either chain can be missing in some cells. Additionally, many experimental designs include a mixture of single cell and bulk heavy chain sequencing. We show that if all heavy chains are paired with a light chain, combining paired heavy and light chain sequences almost always results in more topologically accurate and reproducible B cell phylogenetic trees. However, improvements in topology estimation diminish once ∼30% of B cells lack a light chain, and many tree building methods produce highly biased branch length estimates. Only multi-partition models were able to accurately estimate branch lengths when a significant portion of cells had missing light chains. We conclude by showing that these biases can have a significant impact on conclusions drawn from a study of influenza vaccine response and recommend using multi-partition models when building trees with paired heavy and light chains. We implement all of these methods as part of the Immcantation framework in the R package Dowser v2.0.0, which provides a unified framework for building, visualizing, and analyzing B cell lineage trees.

B cell phylogenetics has a long history and multiple B cell specific methods for tree inference have been developed^14,37–39^. Benchmarking studies of these and more general phylogenetic methods have shown mixed results, possibly due to differences in simulation strategies^16,27,40^. However, previous methods and benchmarking studies focused on heavy chain BCR sequences. Here, we show that including light chain sequences improves tree accuracy, often more than the choice in tree building method. Indeed, on average, all methods tested using paired heavy and light chain sequences had more accurately estimated tree topologies than any method using only heavy chains.

While including light chains almost always improved tree inference when all heavy chain sequences had a paired light chain, most methods had significant biases when some light chains were missing, such as when using a mixture of bulk and single cell BCR data. These biases were most notable in branch length estimates, which represent the estimated number of mutations per site. When some heavy chains did not have paired light chains, maximum parsimony methods underestimated branch lengths, single-partition maximum likelihood methods over-estimated branch lengths, and multi-partition maximum likelihood methods accurately estimated branch lengths (**Figs. 3 and 4**). While all models performed competitively when both heavy and light chain sequences were present for all cells, light-chain dropout can occur in any single cell dataset, leaving some heavy chains without light chains. Because of the possibility of light chain dropout, we recommend multi-partition maximum likelihood methods whenever building trees with paired heavy and light chain sequences, especially if using a mixture of single cell and bulk BCR data.

Multi-partition models are a class of phylogenetic models in which separate parameters, branch lengths, and even topologies can be estimated for different sequence partitions. In this study, we only considered scaled models, in which heavy and light chain partitions use a common tree topology, but all branches are multiplied by a partition-specific scaling factor when performing likelihood calculations. These scaling factors are estimated as free parameters for each partition, allowing light chain genes to mutate at a different rate than heavy chains. These models were chosen because they fit the known biology of BCR evolution: B cells reproduce asexually, so heavy and light chain genes must follow the same topology, and light chains accumulate somatic hypermutations at roughly half the rate of their paired heavy chains (**Fig. 1**). Because only one branch length scalar is estimated for each partition, it is assumed that there is no variation in the rate of heavy vs light chain evolution within a clone^14,15^. Other models exist, such as “unlinked” partition models in RAxML, in which all branch lengths are estimated separately for each partition^15^. Initial experiments with these models found poor performance, especially when some light chains were missing (data not shown). Indeed, the principal benefit of scaled multi-partition models seems to come from assuming limited rate heterogeneity: information from cells with both heavy and light chains helps estimate branch lengths when some cells are missing light chains. This would be lost in fully unlinked models. Future work may incorporate models with relaxed, rather than strict, rate variation among heavy and light chains.

There are some caveats to the results we have presented. The evaluation metrics used in this study – RF cluster distance from the true tree and mean bootstrap score – were limited. RF distance can lack sensitivity and rapidly saturate^41^. However, because we only compare RF cluster distances between estimated and true topologies, we believe these issues do not significantly impact our results. Finally, we only tested one simulation paradigm and set of parameters. However, we argue that biologically realistic simulations are more useful than exhaustive testing of parameter space. A similar framework has been used in a previous (heavy chain) benchmarking study, and we confirmed our simulations produce BCR sequences with realistic heavy and light chain SHM patterns (**Fig. 1**)^16^. Another caveat is that all analyses presented here assume that B cell clones are perfectly identified, and that all V(D)J recombination events occurred before the onset of SHM. Clonal clustering using both heavy and light chains has been addressed in other studies, so in this study we focused on tree building^7,16^.

In summary, incorporating paired light chain sequences into B cell phylogenetic tree building significantly improves both tree accuracy and reproducibility. We recommend using multi-partition maximum likelihood models for building trees with both chains as standard practice for analyzing single cell BCR sequence datasets, especially if they are paired with bulk BCR sequencing. All methods used in this manuscript for building trees with paired heavy and light chains, as well as just heavy chains, are available in the R package Dowser v2.0.0, which is part of the Immcantation suite (immcantation.org)^17^.

## Acknowledgments

This work was funded in part by the National Institute of Allergy and Infectious Diseases grants R01AI104739 (S.H.K.) and K99AI159302 (K.B.H.) and the National Library of Medicine grant T15LM007056 (C.G.J., J.A.S). **Competing interests:** S.H.K. receives consulting fees from Peraton. K.B.H. receives consulting fees from Prellis Biologics. The remaining authors have no competing interests.

## Data availability

All data used are publicly available as part of their respective publications. Simulated data are available for download from Zenodo: https://zenodo.org/record/8338674.

## Code availability

Dowser is an R package for performing phylogenetic analysis on B cell receptor repertoires. All tree building methods are implemented in Dowser, available at https://dowser.readthedocs.io.

The source code and detailed documentation are available on Bitbucket at https://bitbucket.org/kleinstein/projects/src/master/Jensen2023.

---

**Supplementary Figure 1.**
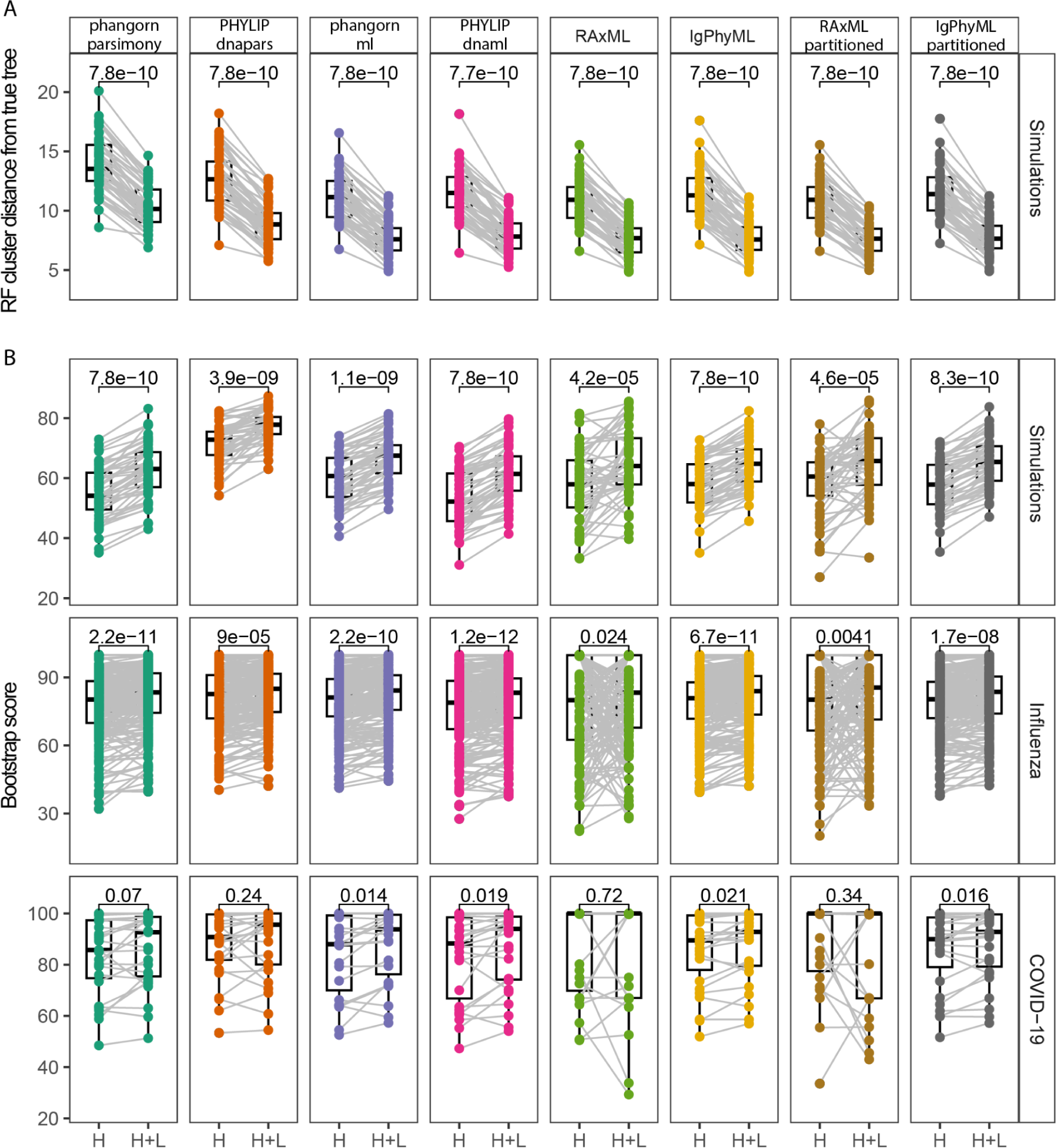
The accuracy and reproducibility of tree reconstruction are improved using paired heavy and light chains. Similar to Fig. 1, but showing boxplots for all eight methods tested. A) Robinson-Foulds (RF) cluster distance between estimated and true tree topologies for trees built using only the heavy chain (H) and paired heavy and light chain sequences (H+L). B) Similar, but with the mean bootstrap value for trees in each dataset. P values were calculated using a Wilcoxon test.

**Supplementary Figure 2.**
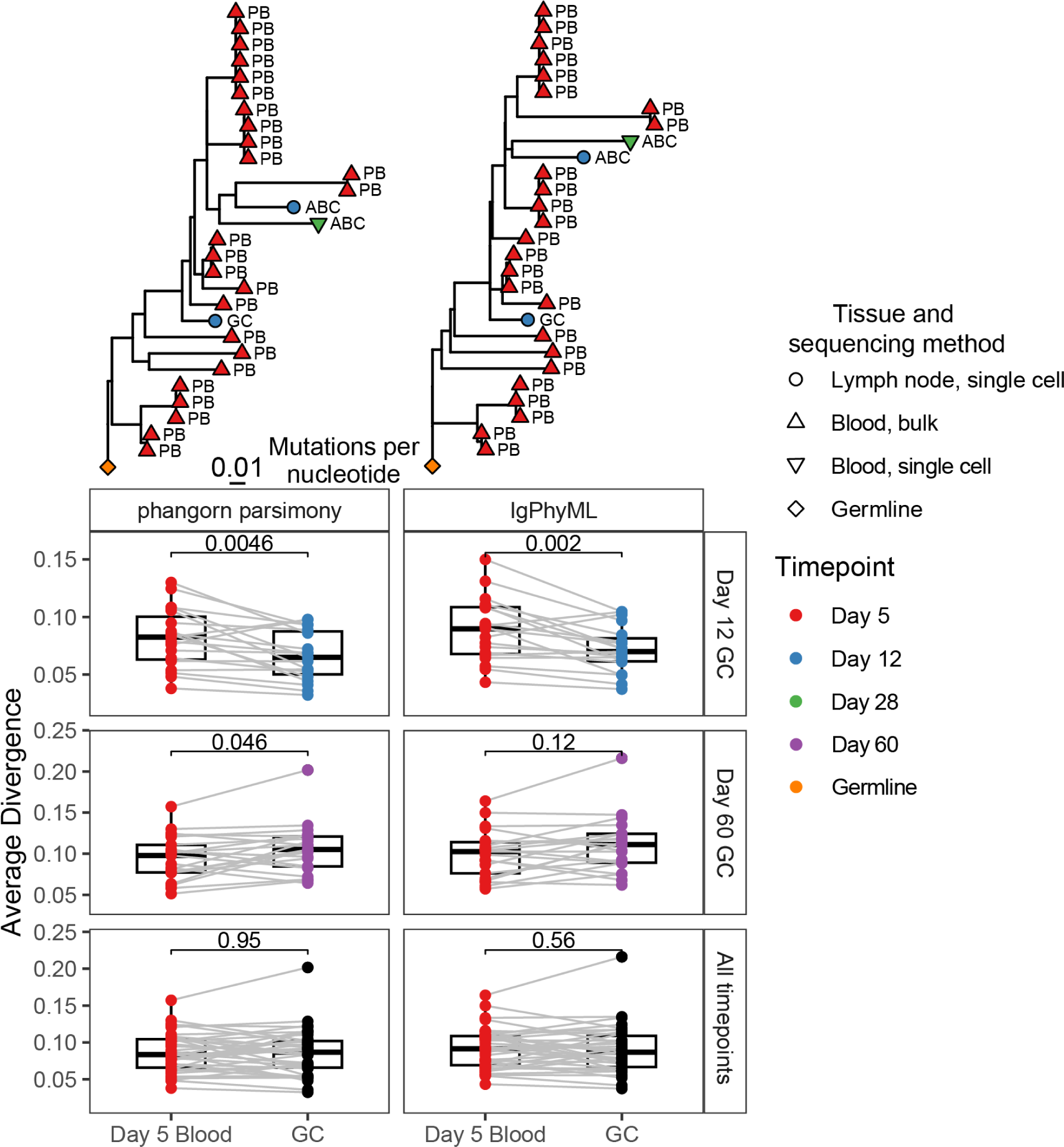
Similar to Fig. 5, but using only heavy chain sequences. Multi-partition maximum likelihood methods were not tested because only heavy chain sequences were included in these comparisons. Above each plot is a representative tree.

